# Evolution of Specialization in Dynamic Fluids

**DOI:** 10.1101/735589

**Authors:** Gurdip Uppal, Dervis Can Vural

**Affiliations:** University of Notre Dame, Notre Dame, IN, United States

## Abstract

Previously we found mechanical factors involving diffusion and fluid shear promote evolution of social behavior in microbial populations ***Uppal and Vural (2018)***. Here, we extend this model to study the evolution of specialization using realistic physical simulations of bacteria that secrete two public goods in a dynamic fluid. Through this first principles approach, we find physical factors such as diffusion, flow patterns, and decay rates are as influential as fitness economics in governing the evolution of community structure, to the extent that when mechanical factors are taken into account, (1) Generalist communities can resist becoming specialists, despite the invasion fitness of specialization (2) Generalist and specialists can both resist cheaters despite the invasion fitness of free-riding. (3) Multiple community structures can coexist despite the opposing force of competitive exclusion. Our results emphasize the role of spatial assortment and physical forces on niche partitioning and the evolution of diverse community structures.

## Introduction

From subcellular structures to ecological communities, life is organized in compartments and modules performing specific tasks. Organelles ***Siegel (1960); Kutschera and Niklas (2005)***, single ***Lewis (2007)*** and multi-phenotype ***Koufopanou (1994); Fu et al. (2018)*** bacterial populations, tissues and organs in multicellular organisms ***Carroll (2001); Hedges et al. (2004)***, casts and social classes in colonial animals ***Beshers and Fewell(2001); Smith et al. (2008)***, and guilds in ecological communities ***Terborgh (1986); Futuyma and Moreno (1988); May and Seger (1986)***, all fulfill specialized roles that are vital for the functioning of the bigger whole. The evolution of specialization also gives rise to cooperative metabolic interdependencies in microbial populations and can serve as a strong mechanism for community assembly ***Zelezniak et al. (2015)***.

There are two classes of evolutionary forces moving a population from having one type of individual performing multiple functions -generalism-, towards one that has multiple types of individuals performing distinct functions -specialism-. The first is “incompatible optimas” ***Solari et al. (2013); Sriswasdi et al. (2017); Goldsby et al. (2012):*** If a population must optimize two functions at once, but if the two phenotypes optimizing these are incompatible, then the population will split into two phenotypes. For example, the somatic and germ cells in volvox colonies are optimized for reproduction and motility. As a result, they have entirely different positioning ***Solari et al. (2006b)***, morphology ***Kirk (2001)***, and protein expression ***Kirk and Kirk (1983)***. In multicellular cyanobacteria, cells differentiate into carbon-fixating cells and nitrogen-fixating heterocysts ***Rossetti et al. (2010)***. E. coli can differentiate into transient non-growing cells and normally growing cells to hedge their bets across different environments ***Lewis (2007)***. A traveling band of E. coli will exhibit a continuum of navigation styles, each specializing in processing different local conditions while still moving in unison ***Fu et al. (2018)***.

A second type of evolutionary pressure originates from the economies of scale. Undertaking one process at high volume is more cost-effective than undertaking multiple processes at low volume. The morphological characteristics necessary to accomplish two distinct functions require two investments in overhead. Specialization is then favored if fitness returns are accelerated by further investment into a specif c task ***West et al. (2015); Cooper and West (2018)***.

In a cooperating community, individuals that “cheat” by diverting resources away from public interest towards self-interest will increase in numbers and eventually dominate the group, leading to a non-cooperative, unfit group. This well-studied evolutionary instability is known as “tragedy of commons” ***Hardin (1968); Rankin et al. (2007); Dionisio and Gordo (2006); Smith (2012)***. From an evolutionary game theory perspective, a specialized community could be viewed as a collection of “semi-cheaters”, in the sense that every sub-population secretes only a subset of the essential public goods but relies on others for the rest. Thus, from game theoretical considerations alone, one might expect a similar tragedy-of-commons scenario, where specialists always eventually dominate a population of generalists. How then should we explain the persistence of generalists in nature, and even the coexistence of various combinations of generalists, specialists and cheaters within one niche?

Evolution of specialization is typically studied in terms of fitness trade-offs or economic considerations. Specialization is shown to occur if relatedness is high and if fitness returns accelerate ***Michod (2007); Michod et al. (2006); Willensdorfer (2009)***; ***Tannenbaum (2007); Rueffler et al. (2012); Taylor (1992); Vural et al. (2015)***. The role of conflicts within groups and spatial assortment is unknown.

Computational models that do consider spatial structure or finite group size, only do so phenomenologically, abstracting away the underlying physics ***Cooper and West (2018); Gavrilets (2010); Willensdorfer (2008); Rueffler et al. (2012); Ispolatov et al. (2012); Menon and Korolev (2015); Gavrilets (2010); Willensdorfer (2008)***. While conceptually useful, such models do not concern with how mechanical forces influence population dynamics and evolution.

Real-life microbial cooperation is mediated by secretions that diffuse and flow ***West et al. (2007)***. Extracellular enzymes digest food ***Greig and Travisano (2004); Bachmann et al. (2011); Pirhonen et al. (1993)***, surfactants aid motility ***Kearns (2010); Xavier et al. (2011)***, chelators scavenge metals ***Griffin et al. (2004); Guerinot (1994); Ratledge and Dover (2000)***; ***JB (1984); Harrison F (2009)***; ***Kümmerli R (2010),*** toxins fight competitors and antagonists ***Mazzola et al. (1992); Moons et al. (2005, 2006)***; ***An et al. (2006); Inglis et al. (2009)***, virulence factors exploit a host ***Zhu et al. (2002); Allen et al. (2016); Sandoz et al. (2007); Kohler T (2009)***, and extracellular polymeric substances provide sheltering ***Mah and O’toole (2001); Xue et al. (2012); Davies (2003)***. Since cells must be within a certain distance to exchange such services, spatial aggregation is often viewed as a prerequisite for multicellular specialization. Spatial effects are known to have a major impact ***Durrett and Levin (1994); Wilson et al. (2003); Fletcher and Doebeli (2009); Wakano et al. (2009)***, and multiple factors can couple together to influence the evolution of cooperation in unexpected ways ***Dobay et al. (2014)***.

In this study, we find that mechanical factors such as diffusion lengths, molecular decay constants and fluid shear forces play an important role in shaping the interaction structure of the community. We find, through first-principles computer simulations and consistent analytical formulas, that microbes self-aggregate to form communities with spatial structure. Contrary to game theoretic intuition, we find that generalist communities can resist specialization. More remarkably, under suitable conditions, we observe that multiple types of community structures can coexist within the same fluid niche. We also determine what physical properties make “socially inhabitable” niches, where free-riders emerge, exploit and invariably destroy both generalist and specialist communities.

Any model aiming to describe evolution of functional specialization must include at least two functions, so that sub-populations can potentially specialize to perform one function each. In the present model microbes can secrete two public goods and a waste/toxin. These molecules diffuse, flow, and decay (cf. ***Figure 1***). The specif c assumptions of our model can be enumerated as follows: **(1)** The system consists of microbes that can secrete two kinds of public goods. A public good refers to a secretion that slows down the growth rate of the producer, but enhances others nearby. **(2)** Every microbe secretes a waste molecule that curbs the growth of those nearby. **(3)** The secretions and bacteria obey the physical laws of fluid dynamics and diffusion. **(4)** Whether a microbe secretes both, one, or none of the public goods is hereditary, except for random mutations. However, every phenotype emits waste.

**Figure 1.**
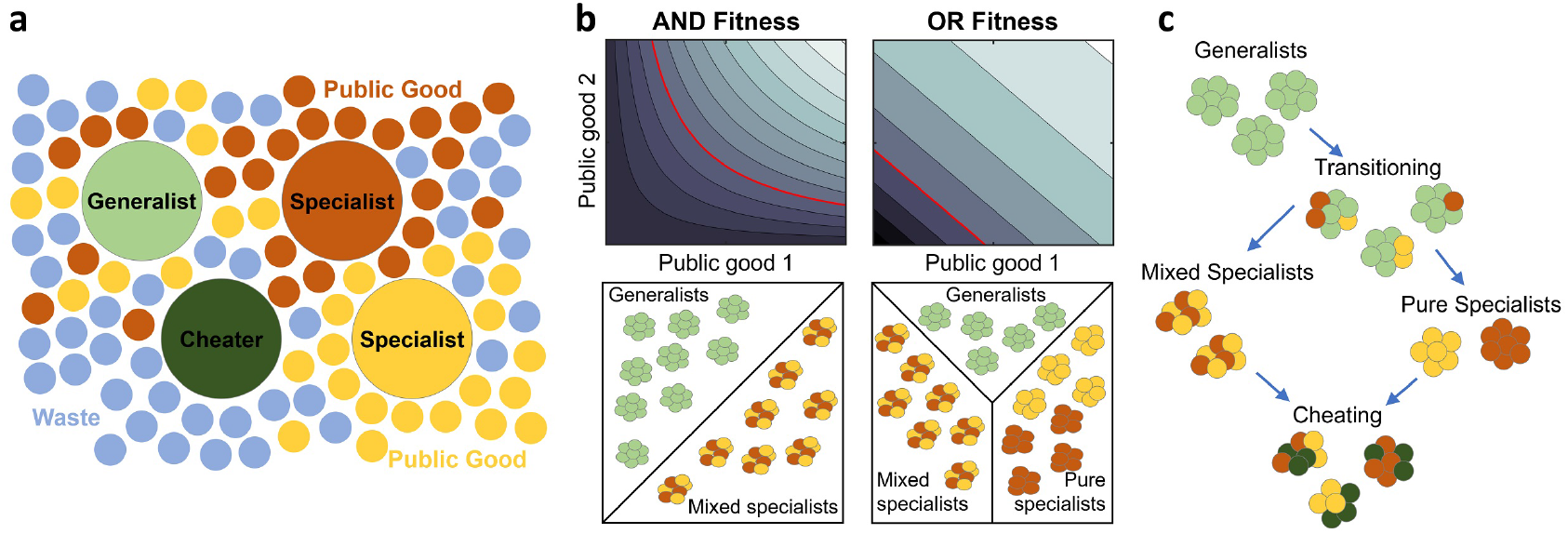
Schematics and dynamics of microbe model. **(a)** Microbes cooperate by secreting public goods into their environment. Generalists (large green circles) secrete two public goods (small yellow and red circles). Specialists (large red and yellow circles) secrete only one of the two public goods. Cheaters (dark green) secrete none of the public goods. All microbes secrete a metabolic waste (small blue circles). **(b)** Fitness contour plots and types of stable groups in each fitness variant. In the top row, we plot the fitness contours as a function of public good concentrations *c*_1_ and *c*_2_. In the bottom row, we show the types of stable groups in each fitness form. The red line represents the contour corresponding to zero fitness. In the AND case, the red line can never cross the *c*_1_, *c*_2_ axes, the fitness is negative when either chemical is not present. Therefore the only types of stable groups are generalists or mixed specialists, as shown in the bottom-left panel. In the OR case, the contours form straight lines. A decrease in one chemical is equally compensated by an increase in the other chemical. The zero contour always crosses the *c*_1_, *c*_2_ axes. It is therefore also possible to have pure specialist groups in the OR case, as shown in the bottom-right panel. **(c)** Evolutionary paths between group types. Groups typically move towards less secretion since cheaters have higher invasion fitness than specialists, who, being “half-cheaters”, have higher invasion fitness than generalists.

We study two models separately. **(5)** In one, which we call AND, access to both kinds of goods is necessary. In the other, which we call OR, both goods contribute to fitness, but the lack of one can be compensated with the other.

## Methods

Our work consists of (1) discrete, stochastic agent based simulations and (2) related continuous deterministic equations. In addition, to gain better analytical understanding, we construct (3) a simple effective model that captures the essential details of (1) and (2).

### Continuous Deterministic Equations

We construct equations governing the number density of three phenotypes *n*_1_(*x,t*), *n*_2_(*x,t*), *n*_3_(*x,t*) two chemical secretions that are public goods *c*_1_(*x,t*),*c*_2_(*x,t*), and a waste compound *c*_3_(*x,t*), as a function of space *x* and time *t*. *n*_3_(*x, t*) is the number density of microbes that secrete both kinds of public goods, to which we refer as “generalists”. The microbes that secrete only public good number one or two, are denoted by *n*_1_(*x, t*) and *n*_2_(*x, t*), to which we refer as “specialists”. Those that secrete no public goods are denoted by *n*_0_(*x, t*), to which we refer as “cheaters”.

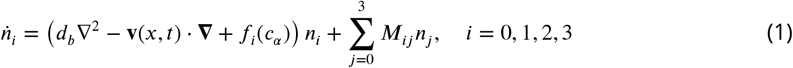

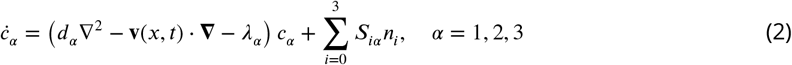

Here indices *i,j* = 0,1,2,3 label phenotypes, whereas the index *α* = 1,2,3 labels chemicals, i.e. the two public goods and waste. Thus, ***Equation 1*** and ***Equation 2*** comprise 7 coupled spatiotemporal equations.

In both equations, the f rst two terms describe diffusion and advection. The flow field **v**(*x,t*) is a vector valued function of space and time, and includes all information pertaining the flow patterns in the environment. In general, it is obtained by solving separate fluid dynamics equations. Mutations between phenotypes and secretions of each phenotype is governed by two matrices

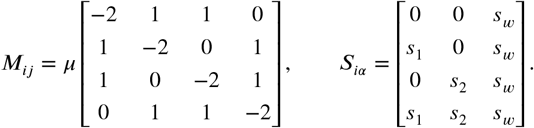

The secretion rate of chemical *α* by phenotype *i* is given by the matrix element ***S**_iα_*, and its decay rate by *λ_α_*. The mutation rate from phenotype *j* to *i* is given by ***M**_ij_*. The diagonal elements ***M**_ii_* indicate the rate at which phenotype *i* mutates into a different category.

In our model, the secretion of public goods is binary, i.e. a good is either secreted or not. Mutations toggle on and off with probability *μ* whether an individual secretes either public good. A mutation can cause a generalist to become a specialist, but two mutations, one for each secretion function, are required for a generalist to become a cheater. Same with back mutations.

The fitness function *f_i_*(*c_α_*) determines the growth rate of phenotype *i*. We consider two cases separately: when both public goods are necessary for survival (AND) and when the public goods can substitute for one other (OR).

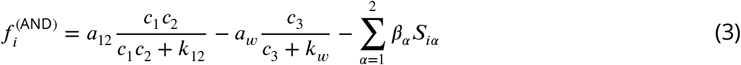

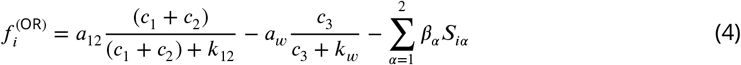

As we see, in both cases, growth rate increases with the local concentration of public goods, *c*_1_,*c*_2_ and decreases with the concentration of waste, *c*_3_. *β_α_* is the cost of secreting public good *α*, so that growth of phenotype *i* is curbed by an amount proportional to its public good secretion. Note that waste is produced without any cost.

Note that with increasing concentration of goods, microbes receive diminishing returns. Similarly, with larger waste, death rate approaches a maximum value. These functional forms are well understood, experimentally verified, and commonly used in population dynamics models ***Monod (1949)***. *a’s* and *k’s* are constants defining the initial slope and saturation values of growth and death.

### Discrete Stochastic Simulations

Our analytical conclusions (cf. ***Appendix 1***) have been guided and supplemented by agent based stochastic simulations. Videos of these simulations are provided in supplementary videos. Our simulation algorithm is as follows: at each time interval, Δ*t*, the microbes (1) diffuse by a random walk with step size *δ* derived from the diffusion constant and a bias dependent on the flow velocity, 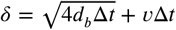, (2) secrete chemicals locally that then diffuse using a finite difference scheme, and (3) reproduce or die with a probability dependent on their local fitness given by *f*(*c_α_*)Δ*t*. If *f* Δ*t* is negative, the microbes die with probability 1, if *f* Δ*t* is between 0 and 1 they reproduce an identical offspring with probability *f* Δ*t*. Upon reproduction, random mutations may alter the secretion rate of either public good -and thus the reproduction rate-of the microbes. Mutations occur on each secretion function with probability *μ* and turn the secretion of the public good on or off. The secretion rate is assumed to be heritable, and constant in time. Numerical simulations for figures were performed by implementing the model described above using the Matlab programming language and simulated using Matlab (Mathworks, Inc.). The source code for discrete simulations is provided as a supplemental f le. A summary of the system parameters is given in ***Table 1***, along with typical ranges for their values used in the simulations. The relevant ratios of parameters are consistent with those observed experimentally ***Kim (1996); Ma et al. (2005)***. Note also that the choice of parameters will be restricted to ensure a finite stable solution is possible. For example, we enforce the quantity *α*_12_ – *α_w_* – 2*βs*_12_ < 0. This is because, if this quantity were positive, then a dense population where the Hill terms in the fitness functions are saturated, will continue to have a positive fitness and grow indefinitely. In the case where secretion rate and/or production cost are low, the waste term is crucial to ensure a finite carrying capacity. We therefore choose *α_w_* ≥ *α*_12_. Other constraints on existence and stability are derived in our Turing analysis (see ***Appendix 1***).

**Table 1:** Summary of system parameters.

| Parameter | Definition | Values for OR | Values for AND |
| --- | --- | --- | --- |
| $d_b$ | Microbial diffusion constant | $0.3906 \times 10^{-6} \text{ cm}^2 \text{ s}^{-1}$ | $1.0 \times 10^{-6} \text{ cm}^2 \text{ s}^{-1}$ |
| $d_1$ | Public good 1 diffusion constant | $5 \times 10^{-6} \text{ cm}^2 \text{ s}^{-1}$ | $5 \times 10^{-6} \text{ cm}^2 \text{ s}^{-1}$ |
| $d_2$ | Public good 2 diffusion constant | $5 \times 10^{-6} \text{ cm}^2 \text{ s}^{-1}$ | $5 \times 10^{-6} \text{ cm}^2 \text{ s}^{-1}$ |
| $d_w$ | Waste diffusion constant | $15 \times 10^{-6} \text{ cm}^2 \text{ s}^{-1}$ | $15 \times 10^{-6} \text{ cm}^2 \text{ s}^{-1}$ |
| $\lambda_1$ | Public good 1 decay constant | $5.0 \times 10^{-3} \text{ s}^{-1}$ | $5.0 \times 10^{-3} \text{ s}^{-1}$ |
| $\lambda_2$ | Public good 2 decay constant | $5.0 \times 10^{-3} \text{ s}^{-1}$ | $5.0 \times 10^{-3} \text{ s}^{-1}$ |
| $\lambda_w$ | Waste decay constant | $1.5 \times 10^{-3} \text{ s}^{-1}$ | $1.5 \times 10^{-3} \text{ s}^{-1}$ |
| $k_{12}$ | Public goods saturation | 0.01 | $3 \times 10^{-5}$ |
| $k_w$ | Waste saturation | 0.1 | 0.1 |
| $s_1$ | Public good 1 secretion rate | $5.0 \times 10^{-3} \text{ s}^{-1}$ | $0.01 \text{ s}^{-1}$ |
| $s_2$ | Public good 2 secretion rate | $5.0 \times 10^{-3} \text{ s}^{-1}$ | $0.01 \text{ s}^{-1}$ |
| $s_w$ | Waste secretion rate | $0.01 \text{ s}^{-1}$ | $0.09 \text{ s}^{-1}$ |
| $a_{12}$ | Benefit of public goods | $7.5 \times 10^{-3} \text{ s}^{-1}$ | $6.5 \times 10^{-3} \text{ s}^{-1}$ |
| $a_w$ | Harm of waste compound | $8.0 \times 10^{-3} \text{ s}^{-1}$ | $10.5 \times 10^{-3} \text{ s}^{-1}$ |
| $\beta_1$ | Cost of secretion 1 | 0.01 to 0.26 | 0.01 to 0.15 |
| $\beta_2$ | Cost of secretion 2 | 0.01 to 0.26 | 0.01 to 0.15 |
| $\mu$ | Mutation rate | $5.0 \times 10^{-8} \text{ s}^{-1}$ | $2.0 \times 10^{-7} \text{ s}^{-1}$ |

### Simple Effective Model

To gain better analytical understanding, we supplement our numerical results with an effective model given in ***Appendix 2***. Our effective model is based on the observation that microbes aggregate into self-reproducing cooperative groups. Different group types, rather than phenotypes, constitute the basic building blocks of our effective model; and the fragmentation rates of these group types constitute the basic parameters of the model. These parameters are “measured” from simulations and depend on the physical properties of the system. The results of our effective model are compared to simulation results in ***Figure 3***.

## Results

### Cooperative groups as Turing patterns

Through numerical simulations and analytical formulas, we see that the system gives rise to spatially segregated cooperating groups in a certain parameter range, as shown in ***Figure 2***. Spots or stripes in reaction diffusion systems are known as Turing patterns, which form whenever an inhibiting agent diffuses faster than an activating agent. In our model the inhibiting and activating agents are the waste and the public goods.

**Figure 2.**
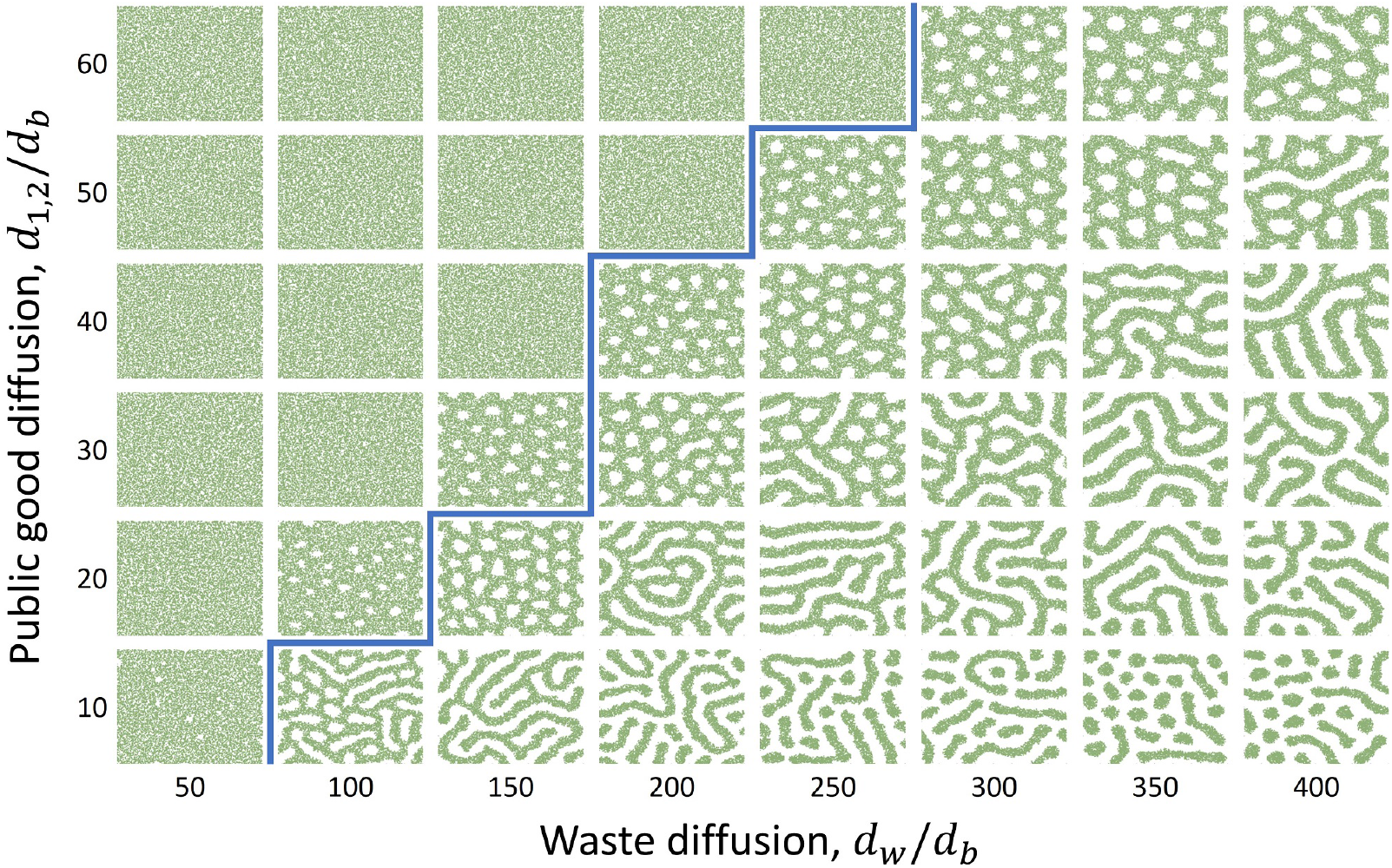
Cooperative groups as Turing patterns. In both the AND as well as the OR fitness type, microbes form aggregate structures via a Turing instability. Here we show generalists for the AND case, similar results hold for the OR case. The thick blue line gives the theoretical region of Turing instability, which agrees with computer simulations see ***Appendix 1***. The population can be homogeneous or form stripes or spots. These patterns can also grow and fragment, forming new colonies. The diffusion constants are normalized by the bacterial diffusion constant, *d_b_* = 1 × 10^−6^ cm^2^s^−1^. Secretion constants used are *s*_1_ = *s*_2_ = 0.015 s^−1^ and *s_w_* = 0.005 s^−1^, public good cost is *β* = 0.1. The rest of the parameters are as given in ***Table 1***.

In general, the structure and size of these cooperating groups will vary with physical parameters. We show in ***Figure 2*** how the pattern forming Turing domain varies with diffusion constants. Our analytical result, derived in ***Appendix 1***, shown by the thick blue line, delineates the parameter space into pattern forming and non-pattern forming regions. While simulations agree well with analytical results, we see some patterns slightly beyond the theoretical region. This is due to the stochastic nature of the simulations which is known to widen the pattern forming region ***Butler and Goldenfeld (2009); Biancalani et al. (2010)***.

In our simulations, we observe that cooperative groups of microbes, i.e. spots and stripes, grow and fragment, thereby giving rise to new structures of the same type. These spatial structure of these patterns differ between generalists and specialists, and therefore have a strong effect on the evolutionary trajectory of the system.

### Effects of secretion cost on specialization

We next determine the roles of secretion cost *β_α_*, and flow shear on group structure and hence specialization. To see the effect of trade-offs on specialization, we varied the cost of public good secretion and determined when specialization occurs in both AND and OR fitness forms. To simplify our analysis, we set *s*_1_ = *s*_2_ = *s*. In order for both types of specialists to then coexist, we also set *β*_1_ = *β*_2_ = *β*. Therefore, generalists pay an overall cost of 2*β*, specialists pay *β*, and cheaters pay no cost. As such, a specialist mutant will invade a generalist group, and a cheater mutant will invade a specialist group. In the absence of spatial structure and flow, the entire population will be dominated by cheaters and will go extinct.

What can we say about the competition between different group types (as opposed to between different strains within a group)? Since the fitness benefit per cost diminishes with larger secretion, one might expect that increasing the cost of the goods would favor the specialists over generalists. Counterintuitively, we find the opposite. Specialist groups indeed grow faster and form larger, expansive, and denser groups, which however are at once taken over by mutants. In contrast, generalists form smaller, sparser, weaker groups that fragment often, which limits the spread of mutants. Therefore, at higher cost *β*, the “weak” generalists are able to coexist and even dominate “strong” specialists (***Figure 3a,b***).

**Figure 3.**
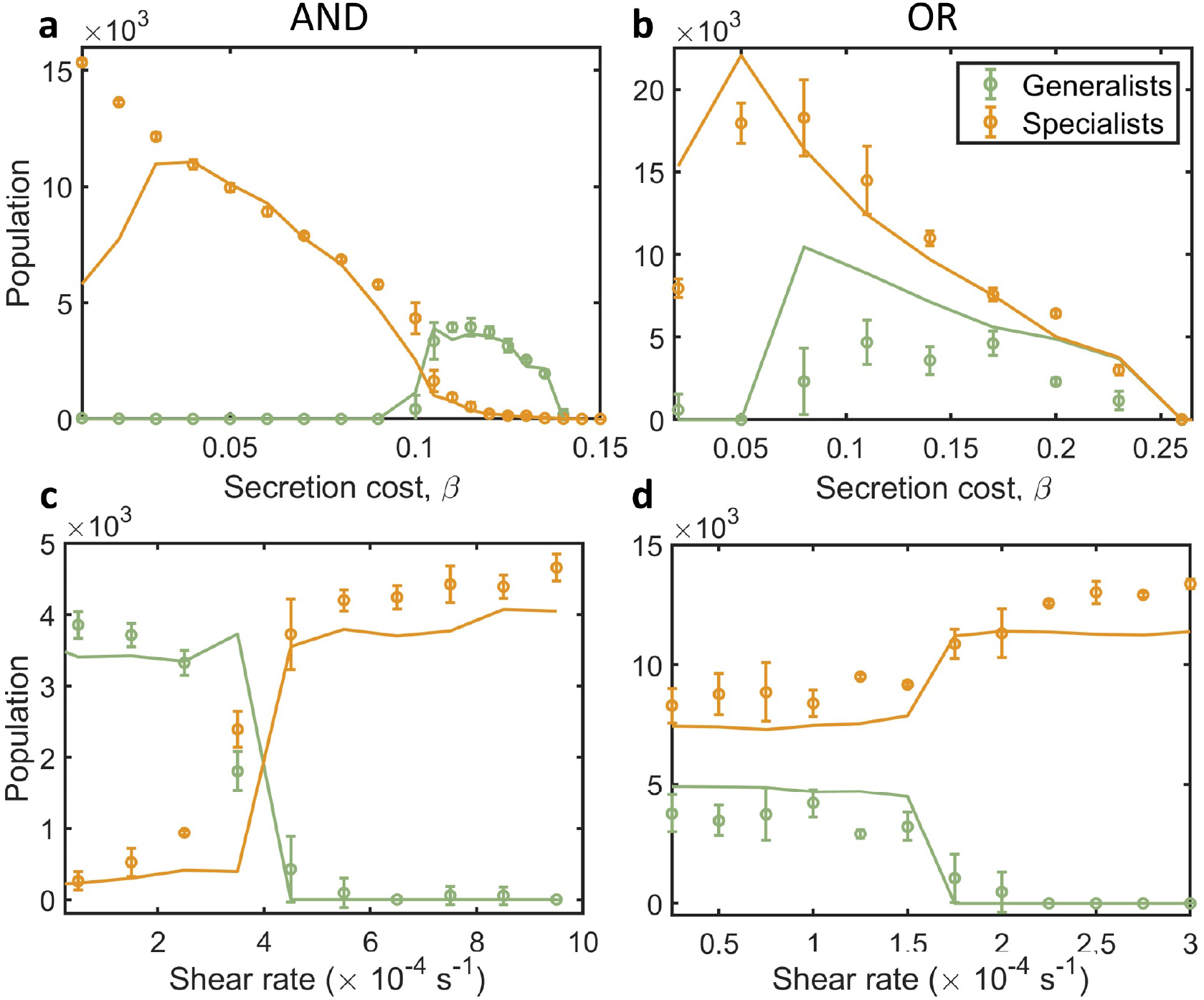
Effects of cooperation cost and fluid shear on specialization. Points with error bars represent numerical results averaged over 5 runs at simulation time *T* = 2 × 10^6^ s in a domain of size 20 mm × 20 mm. Error bars correspond to one standard deviation from the mean. Solid lines are from our effective theoretical model given in ***Appendix2***. (**a,b**) In each fitness variant, we see that specialization and cheating is more abundant for low costs. At lower costs, generalist groups are “too fit” and form large aggregate structures that are more susceptible to mutations. At higher costs, we see generalists out-compete specialists, in the AND case (**a**), and coexist with specialists in the OR case (**b**). At low costs, specialists again dominate. (**c,d**) Effects of fluid shear. A shearing flow causes groups to fragment quicker and stripes to elongate and grow larger. (**c**) In the AND case, with secretion cost *β* = 0.12, we observe that shear transitions the system from a coexisting state to a specialist state. Here, shear causes generalists groups to elongate and become more susceptible to mutations and causes specialist groups to fragment quicker than generalists. (**d**) In the OR case, at cost *β* = 0.17, shear again causes a transition from a coexisting state to a pure specialist state. We therefore see in both cases that flow shear will promote specialization. Parameters not shown in plots are given in ***Table 1***. **Figure 3-video 1**. Video shear stable generalists without flow in AND fitness form. **Figure 3-video 2**. Video shear induced stable specialists in AND fitness form. Same system as video 1, with fluid shear added.

In general, a large uniform population is more susceptible to invading mutants. In contrast, when the population is organized as fragmenting patches, the community structure will prevail as long as the fragmentation rate is larger than the invasive mutation rate. Thus, the type, size, growth and fragmentation of the groups ultimately dictates whether generalism, specialism, or a coexistence of group types are evolutionarily stable.

### Effect of flow patterns on specialization

A shearing fluid flow has been shown to modify social behavior by enhancing the group size and fragmentation rate ***Uppal and Vural (2018)***. We therefore expect the shear rate to effect the types and sizes of groups that we see at steady state.

For constant shear we used a planar Couette flow, with velocity profile and shear rate given as,

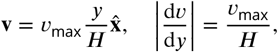

where *ν*_max_ is the maximum flow rate and ***H*** is the height of the domain. We used periodic boundary conditions along the left and right walls (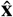 direction), and Neumann boundary conditions for the top and bottom surfaces (*ŷ* direction).

The effect of shear is in general non-trivial and will depend on the group structure observed. We find that a shearing flow enhances group fragmentation rate when a species forms distinct spot patterns and enlarges aggregates when they are closer to forming stripes.

In ***Figure 3c,d*** we show the effect of shear at intermediate costs, where its effect is strongest. We found in both cases that larger shear helps specialists by enhancing their fragmentation rate and enlarging generalist groups. Here, fluid shear transitions the system from a generalist or coexisting state to a specialist state (***Figure 3-video 2***). Thus, fluid shear promotes specialization.

Since advective fluid flow is something that one can tune in an experimental or industrial setting, it is exciting to think of possibilities where flow is used to control the social evolution of a microbial community. Furthermore, since shear is in general spatially dependent, we can use different velocity profiles to localize this control to different regions.

### Combined effects of public good benefit, cooperation cost, and competition on evolution of specialization

We study how varying public good benefit, production cost, and waste diffusion affect the stability of different community structures (***Figure 4***). We find in general, that higher waste diffusion and higher public good benefit helps specialists and higher secretion cost favors generalists. The pie charts also show what conditions leads to coexistence of different group types.

**Figure 4.**
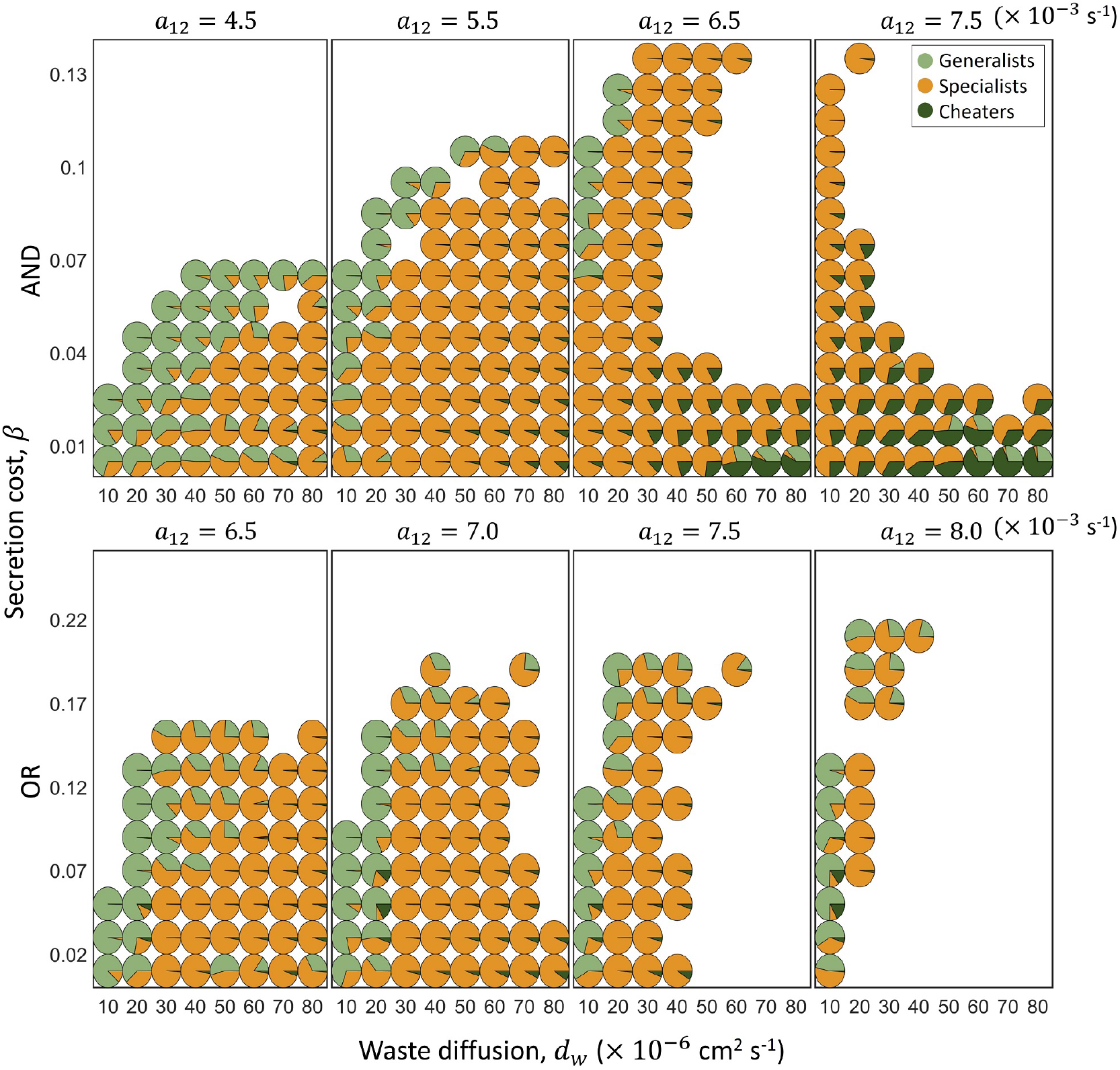
Effects of waste diffusion, public good benefit, and cooperation cost on specialization. Piecharts give the population ratios of generalists, specialists, and cheaters for varying secretion cost *β*, waste diffusion *d_w_*, and public good benefit *α*_12_. Population values were obtained by taking a time average over 1 run for each parameter value. The time average taken over time steps *T* = 1 × 10^6^ s to *T* = 2 × 10^6^ s. When waste diffusion is roughly larger than the public good diffusion, the population can form spatial structures. Under certain conditions, we see coexistence amongst all three – generalists, specialists, and cheaters. At medium costs, cheaters are able to take over faster than groups reproduce, leading to extinction, seen as empty regions. At higher costs, specialists form smaller groups that fragment quicker than cheaters take over and are stable at steady state. Specialists do better overall when waste diffusion is large, since they can then form denser groups without over-polluting themselves. Higher public good benefit, *α*_12_, also helps specialization, since secreting fewer public goods still gives a large benefit. Interestingly, higher benefit also leads to more extinct states, since cheaters can take over quicker. These trends apply to both AND and OR cases. Parameter values for *α*_12_,*d_w_*, and *β* are shown in the plots. Public good diffusion for the AND case is *d*_12_ = 20 × 10^−6^ cm^2^ s^−1^, for the OR case *d*_12_ = 25 × 10^−6^ cm^2^ s^−1^; the rest of the parameters are as given in ***Table 1***. **Figure 4-video 1**. Video of stable specialists in AND fitness form. **Figure 4-video 2**. Video of stable specialists in OR fitness form. **Figure 4-video 3**. Video of generalist/specialist coexistence in OR fitness form. **Figure 4-video 4**. Video cheaters coexisting in AND fitness form.

If waste diffusion is large, self-competition is lower, and specialists can form denser groups without over-polluting themselves. They can then better utilize public goods secreted by their neighbors. If the public good benefit, *α*_12_, is large, specialists also do better since secreting fewer public goods still gives a large benefit (***Figure 4-video 1, Figure 4-video 2***).

As we have already seen, specialization actually occurs more often when trade-offs are small, i.e. at smaller *β*. At higher *β*, generalists are able to coexist with specialists and constitute the majority of the population (***Figure 4-video 3, Figure 3-video 1)***.

We also see that can cheaters persist stably with the population when their invasion f tness is lower than the growth rate of producers. This occurs in regions where producers do not form groups but grow either as stripes or homogeneously in space, which happens when public good benefit is large and when secretion costs are low. In this case, cheaters “chase after” producers, which grow into free space (***Figure 4-video 4***). High waste diffusion also helps cheaters, since they are able to chase producers without over-polluting themselves or their hosts. When their invasion fitness is about equal to the producer growth rate, cheaters take over fully, driving the population to extinction. When the population aggregates into groups, cheater growth is limited to the group. Cooperation then prevails if groups reproduce faster than cheaters emerge. This happens when secretion costs are large. Higher secretion costs can therefore stabilize specialist populations against cheater invasion.

We see two regions of extinction: when public good benefit and waste diffusion are large, at medium costs; and when public good benefit and waste diffusion are low, at high costs. The first case is due to cheaters taking over groups, leading to the tragedy of the commons. Interestingly, this occurs more with higher public good benefit. The population of producers becomes “too fit” and more vulnerable to cheating mutations. For the second case, since costs are high and benefits are low, microbes need to form dense groups to utilize enough goods to be stable. However, due to the low waste diffusion, these groups over-pollute themselves and are no longer stable.

We see similar trends for both the AND and OR cases. The main distinction between the two being, for the OR case, we predominately see pure specialist groups and only have mixed specialists in the AND case. We do not see many mixed specialists in the OR case since mutations take over generalists groups quicker and stabilize as pure groups, whereas in the AND case, pure groups would die out unless the complementary specialist also evolves in the same group. The AND structure is therefore essential to have true division of labor, where each type of specialist exists equally in the group.

### Localization of specialization and coexistence in pipe and vortex flows

We next study the evolution of specialization in Hagen-Poiseuille and Rankine vortex flows. Again, we set the cost parameter to a value where shear makes the biggest difference. As with the case with constant shear (***Figure 3c,d***), we set for AND fitness, *β* = 0.12 and for OR fitness, *β* = 0.17. For a Hagen-Poiseuille flow in a two-dimensional pipe, the flow rate and shear rate are given by,

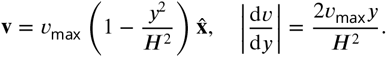

From our results with a constant shear (***Figure 3c,d***), we expect higher shear regions of the pipe to be occupied by specialists and lower shear regions to be occupied by generalists. However, we see the opposite to occur (***Figure 5***). This is due to boundary and second order effects. Generalist groups on the boundary fragment more often, and longer groups are formed in regions of intermediate shear (***Figure 5-video 1***). The fragmenting generalists groups act as a source for specialists groups in the intermediate regions of the pipe. Near the center of the pipe where the shear rate is low, groups do not fragment as quickly and are taken over by cheaters. We therefore see a coexistence of species across the pipe, with generalists at the boundary, followed by specialists in the intermediate regions, and no one at the center (***Figure 5***).

**Figure 5.**
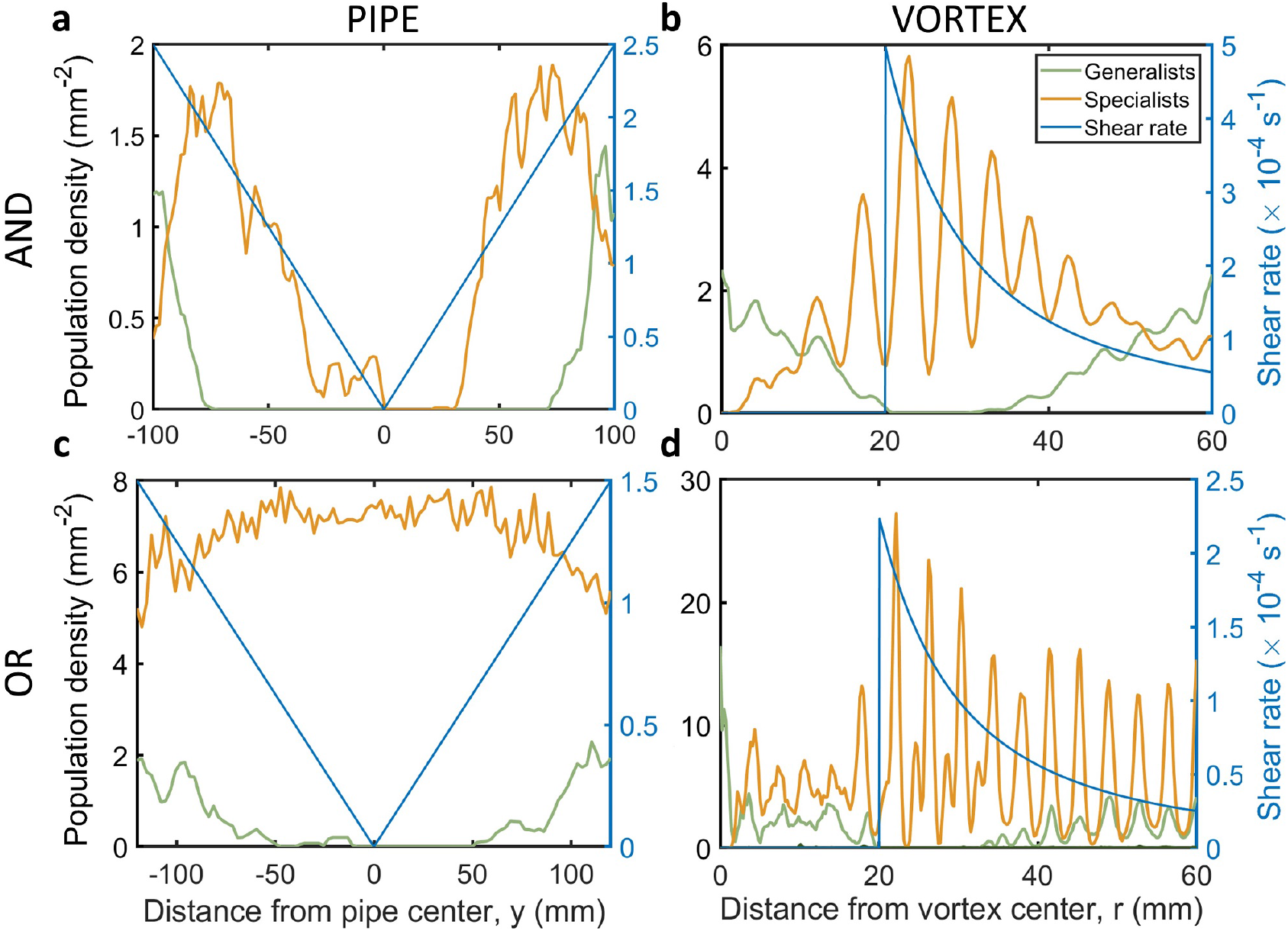
Coexistence of specialists and generalists in pipes and vortices. The local shear profile dictates which species is stable. When shear is spatially varying, we can get coexistence of generalists and specialists. (**a**) For the AND case, in a pipe with Poiseuille flow, we see generalists residing at the boundaries followed by specialists towards the middle. In the center where shear is lowest, cheaters are able to take over and destroy groups, leading to extinction. (**c**) For the OR case, in a pipe, we see coexistence between generalists and specialists at the boundary and specialists predominately in the center. The mutation rates used for the pipe are *μ* = 5 × 10^−7^ s^−1^ for the AND case and *μ* = 5 × 10^−8^ s^−1^ for the OR case. (**b,d**) In a Rankine vortex flow, we see generalists where shear is lowest, and specialists residing in an annulus where shear is at its maximum. The mutation rates used for the vortex are *μ* = 3 × 10^−7^ s^−1^ for the AND case and *μ* = 5 × 10^−8^ s^−1^ for the OR case. Secretion costs used are *β* = 0.12 for the AND fitness, and *β* = 0.17 for the OR fitness. The rest of the parameters are as given in ***Table 1***. Average pipe population densities were obtained from averaging 5 runs in a domain of size 200 mm × 40 mm at a simulation time of *T* = 2 × 10^6^ s. Average vortex population densities were obtained from averaging 5 runs in a domain of size 60 mm× 60 mm also at a simulation time of *T* = 2 × 10^6^ s. **Figure 5-video 1**. Video of evolution and coexistence in pipe for AND fitness. **Figure 5-video 2**. Video of evolution and coexistence in vortex for AND fitness.

Next we study evolution in a Rankine vortex. The flow and shear profiles for a Rankine vortex with radius ***R*** and circulation Γ are given by,

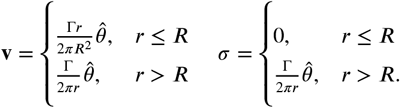

The distribution of specialists and generalists in the vortex agrees better with previous results from constant shear (***Figure 5***). We see generalists persist in regions of low shear and specialists mainly reside in an annular region where shear is large (supplementary video ***Figure 5-video 2***).

In either case we see coexistence of different species across the full domain. The local shear profile dictates which species is stable in each region. A varying shear profile can therefore allow for the coexistence of species across an environment.

#### Box 1. Summary of results

**Specialization:**

- *Large waste diffusion:* Larger waste diffusion lowers self-competition and allows specialists to form denser groups to better utilize public goods secreted by neighbors.
- *Large public good benefit:* A high benefit for public goods allows specialists to still be fit without secreting as many public goods. This also helps cheaters exploit producers.
- *Lower secretion costs:* A lower secretion cost can help specialists dominate over generalists, since a smaller penalty for cooperation can make generalists “too fit”. In this case, large generalist structures are easily taken over by specialist mutants.
- *Group structure:* Specialists from groups when waste diffusion is larger than public good diffusion and when costs are not too low. When specialists do not form groups, they are easily taken over with cheaters, leading to either “chasing cheaters” (***Figure 4-video 4)*** or extinction. When generalists form smaller, reproducing groups, they are able to escape take-over by specialists and out-compete specialists.
- *Fitness type:* The fitness type dictates which types of specialists structure we see - pure or mixed. In the OR case, specialists generally evolve into structures of isolated types of specialists, see ***Figure 4-video 2***. The AND structure is therefore essential to have true division of labor, where each type of specialist exists equally in the group, see for example ***Figure 4-video 1***.

**Cheater coexistence:**

- *Lack of group structure:* Cheaters cannot exist on their own, but must “predate” on producers - generalists or specialists. When producers are fit, and do not form groups, they can grow quicker than cheaters fully taking over. This occurs when waste diffusion is large, and when secretion costs are low.
- *Small invasion fitness:* When cheaters take over slower than producers reproduce, they are able to coexist. This happens when secretion costs are low, since the advantage of not secreting is lower (***Figure 4-video 4)***.

**Extinction:**

- *When cheaters take over:* when the mutation rate and invasion fitness of cheaters large enough such that they take over groups faster than they fragment. This happens when public good benefit is large and waste diffusion is large, we see this in the right-middle regions of plots in ***Figure 4***.
- *When groups are not stable:* When costs are large and public good benefit is low, cooperators need to form denser groups to increase fitness. However, with low waste diffusion, denser groups over-pollute themselves and are no longer stable. We see this in the top-left regions of plots in ***Figure 4***.

**Fluid shear:**

- *Enhanced group fragmentation:* A shearing flow stretches and distorts groups. It can help groups fragment and reproduce quicker, allowing stability over cheating mutations ***Uppal and Vural (2018)***.
- *Enhanced specialization:* Shearing flow can help specialist groups fragment quicker than generalist groups, and therefore transition a population to contain more specialists (***Figure 3-video 2***).
- *Coexistence of group types:* The local shear rate can determine what groups are stable. A spatially varying flow profile can then allow for coexistence of different community structures across the full fluid domain (***Figure 5-video 1, Figure 5-video 2***).

## Discussion

It is well known that spatial structure is key in the evolution of cooperation. By forming reproducing groups, organisms are able to combat takeover by cheating mutants. In a similar vein, spatial structuring can have a large effect on the evolution of specialization.

Previously we had found that mechanical factors involving diffusion and fluid shear can overcome tragedy of the commons ***Uppal and Vural (2018)***. The mechanism gives rise to a variant of Simpsons paradox ***Chuang et al. (2009)*** where individual groups may decrease in sociality, but the population as a whole becomes more social. The present study is also based on the “evolution in a fluid” model in ***Uppal and Vural (2018)***, but with the addition of *multiple* kinds of public goods, to allow for functional specialization and multiple kinds of community structures.

We have studied a physically realistic advection-diffusion system describing microbial growth and evolution. Through this first principles approach, we determined the effects of diffusion, public good benefit and production cost, as well as flow patterns on the evolution of specialization. We showed that specialization is stable when a transition to specialization results in smaller, faster fragmenting groups or when it induces a discrete group structure from a spatially homogeneous generalist state.

In nearly all studies of specialization, the basic mechanism is a trade off between different biological functions. For example, the trade off between mobility and reproduction in volvox ***Solari et al. (2006a, 2013)***. Multicellular collectives are then able to split these tasks amongst different components. We therefore expect that a larger cost in performing both tasks should favor specialization. However, we find that specialization can occur when the trade-off is small, and generalists can persist even when the trade-off is large. In this case, even though a transition to specialization may afford a greater fitness gain, generalists are more evolutionarily stable since they form smaller groups that reproduce quicker. Therefore, fitness gains cannot completely determine when specialization will occur; geometric effects, flow and physical forces can outweigh these fitness gains.

We saw that fluid flow can alter the group structure. A shearing flow can increase group size and group fragmentation rates, and therefore change the evolutionary stability of a community structure. When the shear profile varies over space, as in a Hagen-Poiseuille or Rankine vortex flow, we see a coexistence of generalists and specialists. We therefore view the evolution of division of labor as a mechanical phenomena, where the physics of the media dictates the evolutionary stability of its inhabitants. With this perspective, media can be engineered to control the evolution of biological systems to fit our desired application.

Many authors view undifferentiated multicellularity as a prerequisite for specialization ***Pfeiffer and Bonhoeffer (2003); Gavrilets (2010); Bonner (1998); Rossetti et al. (2010); Michod (2007)***. In the case where generalists form a spatially homogeneous population and specialists form groups, we have seen that a transition to specialization can induce a group structure. These groups may then further specialize and develop individuality and germ-soma specialization. In this case, division of labor can be viewed as a driver of multicellularity, rather than just a consequence. Our findings emphasize the spatial and physical mechanisms for the evolution of multicellularity.

## Acknowledgments

This material is based upon work supported by the Defense Advanced Research Projects Agency under Contract No. HR0011-16-C-0062 and National Science Foundation grant CBET-1805157

## Appendix 1

To understand how each type of species competes evolutionarily with the other, we solve for Turing patterns formed in each state, in the absence of mutations. By performing a linear Turing analysis on each system, we determine the regions in phase space where patterns form.

We study the Turing patterns of stable systems of generalists and specialists in both AND and OR fitness forms.

### Generalist Turing analysis

For generalists, we linearize the following system

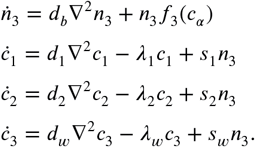

Next, we make the further simplification setting *d*_1_ = *d*_2_ = *d*, *λ*_1_ = *λ*_2_ ≡ *λ* and *s*_1_ = *s*_2_ ≡ *s*, treating both public goods symmetrically. With this simplification, we can set *c*_1_ = *c*_2_ ≡ *c*_12_, and reduce the system to three equations. We then have, for the fitness functions,

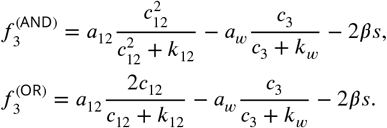

We denote by 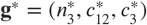 the homogeneous steady state solution, given by setting the reaction terms to zero,

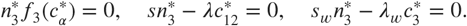

The steady state value for the chemicals are then given by,

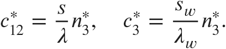

The solution for 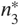 depends on the form of the fitness function. In the AND case, 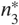 is given by solving the qubic equation,

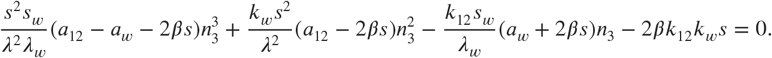

We enforce *a*_12_ – *a_w_* – 2*βs* < 0 for stability, otherwise the dense state will become unstable as the hill forms become saturated. Since the linear and constant coefficients are also negative, by Descartes rule of signs, in order to have any positive solution, we need *a*_12_ – 2*βs* > 0. This implies we have up to two positive solutions, since there are two sign changes.

In the OR case, we have 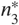 given by solving the quadratic equation,

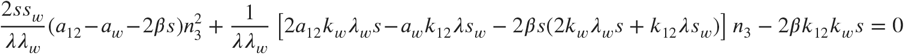

For this to have a positive solution, we require

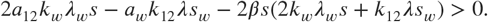

Next we perform a linear stability analysis for each case. We let **g** ≡ (*n*_3_, *c*_12_, *c*_3_)^*T*^ – **g**\*, be a perturbation from the steady state. Our linearized system then looks like,

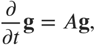

where our stability matrix ***A*** is given as

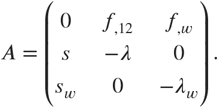

Here *f*_,12_ and *f_,w_*, in each case, are given by

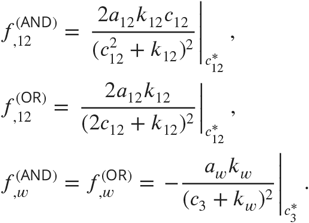

The characteristic polynomial is given in terms of the invariants of a 3 × 3 matrix,

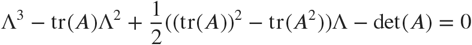

Again, using Descartes rule of signs, we have the following requirements for linear stability, and with the knowledge that tr(***A***) is negative, we require

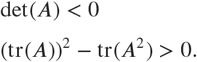

Next we include diffusion and look for a Turing instability. We expand our solution in terms of Fourier modes with exponential growth,

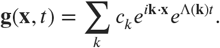

Plugging this into our linearized system, we get the eigenvalue equation, (-*k*^2^*D* + *A*)**g** = Λ**g**. Where,

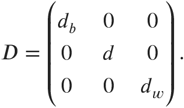

If we denote by ***M***(*k*) = -*k*^2^*D* + *A*, the characteristic equation for this system is now given as

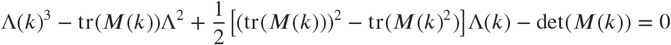

We now use Descartes rule of signs once again, this time to get an instability. First, we have tr(*M*) = -*k*^2^tr(*D*) + tr(*A*) < 0, which makes the quadratic term positive. For the linear term, we have

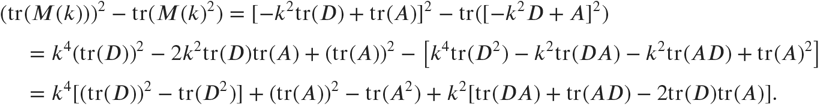

Now since 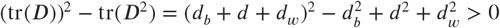, 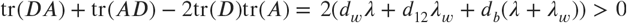, and we require (tr(*A*))^2^ – tr(*A*^2^) > 0 from linear stability, the linear term is also positive. Therefore, by Descartes rule of signs, the requirement for Turing instability reduces to,

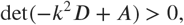

for some range of *k*> 0.

Inspecting this further, we see that det(-*k*^2^*D* + *A*) gives a cubic polynomial in *k*^2^ which goes to negative infinity as *k* → ∞ and passes through det(*A*) < 0 at *k* = 0. We therefore see that a Turing instability begins when the local maximum of Ψ(*k*^2^) = det(-*k*^2^ + *A*) vanishes. We therefore have that the critical wavenumber is given by maximizing Ψ(*k*^2^). Let *w* = *k*^2^, then

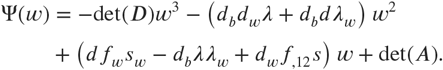

The local maximum is then given by taking 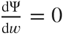 and taking the positive root,

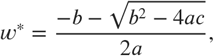

where 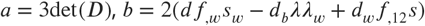, *c* = det(*A*). Given the critical value *w*\* and a public good diffusion *d*, we can find the critical waste diffusion that gives a Turing instability by setting 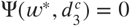 and solving for 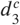. This then gives a relation for 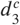 as a function of the other system parameters, when it exists. The theoretical region of Turing instability is compared against numerical results in ***Figure 2***.

### Specialist Turing analysis

The specialist case can be treated in a similar manner. Here we take *n*_3_ = 0 and because of symmetry, take *n*_1_ = *n*_2_ ≡ *n_s_*. We therefore again reduce our system to three equations and get the following system,

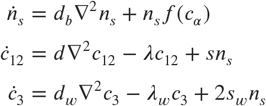

The factor of 2 in the waste chemical term comes from the fact that both populations *n*_1_ and *n*_2_ secrete waste while only one secretes the public good. We also have that the factor of 2 in front of the cost in the fitness functions now becomes a one. We therefore have the same results as in the generalist case with the substitutions,

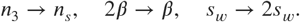

With these substitutions, the results from the generalists case applies also to the specialists. In general, the Turing domain will be different for the two states. These differing Turing regions and pattern types effect what type of state, generalist or specialist, will be more stable evolutionarily when also considering mutations.

## Appendix 2

### Effective group models

To get a better understanding of the results in ***Figure 3***, we developed an effective model of ordinary differential equations describing the dynamics of each type of group. We assume that each type of species, generalists and specialists, form groups that grow and reproduce with different rates. The growth of the groups are given by a logistic equation, with a carrying capacity dependent on the domain. In order to fully describe the logistic growth, we also need to include transient states that are not stable but are continuously formed. The transient states therefore restrict the space in which the stable states are allowed to grow. We denote generalists groups by ***G***. Groups composed of generalists and specialists are transitioning to a specialist state and are denoted by ***T***. Mixed specialist groups made of microbes that secrete different public goods are represented by ***M***. Pure specialists groups composed of microbes secreting only one public good are denoted by ***P***. Pure specialist groups only occur in the OR case, and we exclude them in our effective model for the AND case. Finally, groups with cheaters that secrete no public good are denoted by ***C***. The evolutionary paths the groups can take are illustrated in ***Figure 1c***.

### Effective model for AND type fitness

The group dynamics for the AND case are described by the following set of equations,

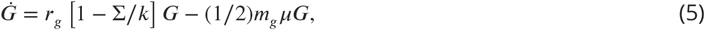

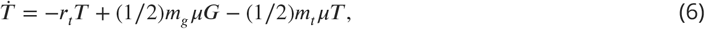

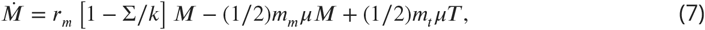

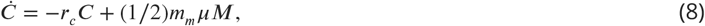

where the constants *m_i_* correspond to the number of microbes in a group of type *i*, and Σ = *m_g_G* + *m_t_T* + *m_m_M* + *m_c_C* is the total population. The rates *r_i_* give the reproduction rates for stable groups and the death rates for transient groups. The carrying capacity for the system is given by *k*. Mutations cause groups to go down in number of secreted compounds (***Figure 1c***), and are given by terms proportional to the mutation rate μ. Back mutations do not stick in groups since they are paying more cost than their neighbors, and so we neglect transitions towards secreting more goods. The factor of 1/2 in front of mutation terms is due to mutations not always sticking in a group before it splits in two. The parameter values were obtained by fitting logistic growth curves to simulations without mutations to determine the natural growth of an isolated species.

By setting (***Equation 5 – Equation 8***) to zero, we obtain the steady state values for each type of group. The possible steady states are the trivial extinct state, where all group populations vanish, a stable state of mixed specialists and cheaters, and a stable state where all species coexist. The specialist/cheater state is given by,

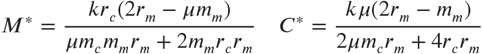

In the stable state where all group types coexist, the number of generalists groups is given by,

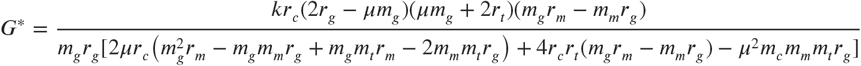

For this to be positive, the generalists group reproduction rate must satisfy,

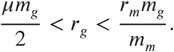

The other species can be given from the ratios,

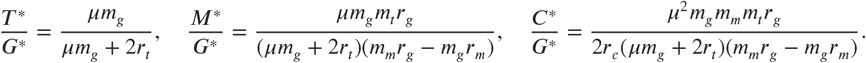

We plot the results of our effective model as solid curves against simulation results (***Figure 3)*** and get good overall agreement with the numerical simulations. At low costs (*β*), microbes form stripes or become homogeneous, and the groups structure assumption of our effective model breaks down.

### Effective model for OR type fitness

In the OR case, pure specialist groups are now stable. Since we treat both chemicals symmetrically, and since the chemicals enter additively in the fitness function, pure specialist and mixed specialist groups are equivalent. However, to form a mixed specialist state, opposite mutations are needed within the lifetime of a generalist group. Since transition states are no longer unstable, once a mutation occurs in a generalist group, it quickly stabilizes to a specialist group. With these considerations, we build and solve an effective model only including generalists, pure specialists, and cheaters. The effective model is described by the following dynamical equations,

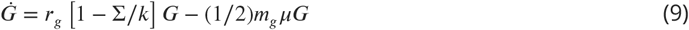

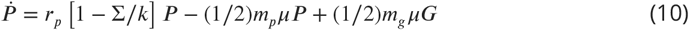

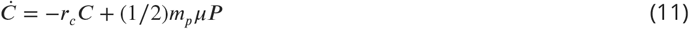

Setting (***Equation 9 – Equation 11***) to zero to get steady states, we again get two non-trivial solutions. One with only specialist and cheating groups, and one where all three coexist. For the solution with only specialists and cheaters, we get

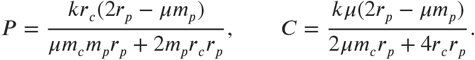

For the solution where all three coexist, we get for the generalists,

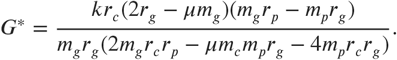

The other groups are given by taking the ratios,

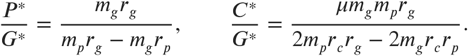

We plot the results of our effective model in the OR case against simulation results (***Figure 3***) and again get a good overall agreement with the numerical simulations.

